# Daily and Weekly Fluctuations in the Microbiome of the Urinary Tract of Postmenopausal Women with No History of Urinary Tract Infections

**DOI:** 10.1101/2025.11.04.686513

**Authors:** Jacob Hogins, Shreya Mekala, Jose Resendiz, Akhil Nelapolu, Sara Papp, Philippe E. Zimmern, Larry Reitzer

## Abstract

At least half of all women will experience a urinary tract infection (UTI) in their lifetime. Age, sexual activity, hormonal status, and the urinary tract microbiome (urobiome) contribute to the probability of infection. While studies have assessed urobiome composition during and after infection, no study has tracked the daily and weekly fluctuations of healthy urobiomes to determine urobiome stability. In this study, we sampled the clean catch urine of 3 women with no history of UTIs over the span of two weeks sampling three times a day on two days each week to track urobiome changes, and whether the urobiome in individuals with no UTI history resembles that of individuals after a UTI. Two participants were dominantly colonized by *Lactobacillus* while the third had a more diverse set of taxa albeit at a lower total amount of bacteria. We previously reported that the urine from only one of the less diverse urobiomes supported the growth of recently isolated uropathogenic *Escherichia coli* or *Enterococcus faecalis*, while the other two participants’ urines did not support growth. We conclude that 1) urobiomes exist in healthy post-menopausal women 2) these urobiomes have a stable and robust community structure and 3) a combination of colonization resistance, including nutrient limitation, and urobiome structure contributes to resistance to UTIs.

**Importance:** Presented here is the first analysis of urinary tract microbiomes (urobiome) of post-menopausal women with no history of urinary tract infections collected at multiple times. These data contribute to the growing understanding of urobiomes of post-menopausal women during a healthy state. Minimal fluctuations were observed on an individual basis, confirming that the time of day or day of the week has no impact on the findings in longitudinal urine sample studies.

## Introduction

Nearly half of all women will experience a urinary tract infection (UTI) in their lifetime and susceptibility increases with age (1). Factors like age, sexual activity, diet, menstruation, menopause, and composition of the urinary microbiome (urobiome) all contribute to a persons’ sus-ceptibility to UTIs (1–3). The existence of the urobiome has been reported since the early 2010s, but these studies have focused primarily on healthy young—pre-menopausal—females or people with urinary abnormalities, cancer, or other comorbidities (3–7). Given that UTI risk increases with age (1) and changes in hormones can affect the urobiome (3), then understanding the affects that menopause and daily factors—e.g., diet—have on the urobiome is as important as understanding the variations in premenopausal females.

A person’s diet impacts nutrient availability in the bladder, which in turn affects how microbes grow and survive in the urinary tract environment (8). The urines utilized in this study have previously been described for their ability to support bacterial growth and were correlated with the diets of the participants during urine collection (9). Briefly, three post-menopausal females with no history of urinary tract infections provided 12 urine samples over a period of two weeks. When inoculated with *Escherichia coli* and *Enterococcus faecalis* only one participant’s urine showed robust bacterial growth while the remaining two supported little growth if any. The growth in relation to diet showed no specific correlation between nutrient intake or time of day with the bacterial growth capacity. Instead, the urines of each individual participant supported different average final densities of bacterial growth (9).

Here we further the investigation into the factors that have protected the participants from UTIs by assessing their urobiomes. We found that these participants not only had a urobiome, but that it contained taxa protective against infections in other body locations—e.g., the vaginal or intestinal tract—which could contribute to the participants never having been infected. The urobiomes were found to be stable over the time course that we analyzed and resembled a urobiome similar to that of a premenopausal female. This comprises the first short-term longitudinal study of urobiomes in healthy post-menopausal participants who have never had a urinary tract infection. Identifying how and whether the urobiome changes and whether healthy naïve bladders contain a microbial presence contributes to our understanding of the factors that influence UTI susceptibility.

## Methods

### Urine Sample Collection

Three post-menopausal females (ages 62, 64, 65) who never had a UTI and were not on hormonal therapy, had diabetes, or had any other urinary tract anomalies or history of kidney stones were selected for this study. Participants provided mid-stream urine samples 3 times daily (8 a.m., 12 p.m., and 4 p.m.), 2 days each week for 2 consecutive weeks for a total of 12 urine samples per participant. Participant A’s first sample was damaged in transfer from the clinic to the lab; thus, patient A only has 11 samples.

Mid-stream urine samples were de-identified after collection and tested for urine pH, specific gravity, glucose, ketones, and leukocyte esterase to ensure the absence of any type of urinary tract infection markers. Samples were frozen at -80°C until transfer from the clinic to the laboratory.

### DNA Isolation and 16S Amplification from Urine Samples

The method was adapted from Pearce et al. 2014 (6). Urine samples were thawed, and 1 milliliter of urine was centrifuged for 10 minutes at 16,000 x g for 10 minutes at 4°C. All the supernatant was discarded, and the pellet was resuspended in 200 microliters of tris (20mM pH 8.0) EDTA (2mM) buffer supplemented with 1.2% Triton X-100, 5 micrograms of lysozyme, and 150U of mutanolysin. Lysis occurred over 90 minutes at 37°C followed by an RNase A digestion for 15 minutes at 37°C and a proteinase K digestion for 20 minutes at 56°C. Lysis was deemed complete once the solution was clear. The protocol for the “DNeasy Blood and Tissue kit” (Qiagen) was followed after completion of the protein digestion. The elution was carried out using 50 microliters of AE buffer (Qiagen) heated to 56°C. Fifteen nanograms of isolated DNA, as determined by nanodrop, was subjected to 16S sequencing following the Illumina 16S sequencing workflow (10) with the following alterations to the first (16S PCR amplification) PCR step: 3 minutes at 95°C; then 30 cycles of 30 seconds at 95°C, 30 seconds at 55°, and 90 seconds at 72°C.

### 16S Sequencing and Data Analysis

To index the isolated DNA, the Illumina workflow was followed exactly. Indexed DNA was sequenced by the Genome Center at the University of Texas at Dallas. Raw reads were processed through the DADA2 (v1.36.0) pipeline (11) and bacterial taxa counts were generated using a Baysian-based least common ancestor (BLCA) taxonomic classification method (BLCA v2.2) (12). Data was assessed using phyloseq (v1.52.0) (13), collapsed to the Genus rank. Relative abundances were generated, followed by UMAP (uwot (v0.2.3)) (14) analysis of CLR transformed features (compositions (v2.0-9)). To calculate beta diversity, vegan (v2.7-2) (15) was used and the data was visualized using pheatmap (v1.0.13) (16). All other graphs were generated with ggplot2 (v4.0.0) (17).

## RESULTS

### Overall Comparative Analysis

Here we assessed the compositions of, and fluctuations in the urobiomes of healthy participants defined by sequencing the V3-V4 region of the 16S ribosomal RNA. When comparing each of the urobiomes, we found 10 (participant A), 105 (participant B), and 88 (participant C) taxa in each urobiome (Supplemental Fig 1) which is within the range seen by other groups who study urobiomes of patients with urinary conditions (6). When comparing each of the biome’s centered log ratio-transformed feature counts, participants B and C both had more similar distributions to one another than participant A (Fig 1). To determine the richness and evenness of the urobiomes, we generated alpha diversity statistics (Shannon and Simpon indexes). Regardless of the participant, there was very little change in alpha diversity throughout the day (Fig 2). Participant C showed a gradual increase in alpha diversity as the day progressed, then returned to a lower alpha diversity each morning. However, these changes were not found to be statistically significant. Participants B and C both had low diversity values due to the overabundance of *Lactobacillus* in both participants, while participant A had a more even distribution of Genera (Fig 3). To determine the differences in the compositions between the participants, we calculated beta diversity statistics (via the “bray” method). A similar pattern to the alpha diversity was observed with these beta diversity measures, where participants B and C were nearly overlapping, but participant A sits on its own (Fig 4A). To remove the skew set by participant A, we regenerated the beta diversity statistic for participants B and C alone. The resulting analysis shows that both participants’ urobiomes are similar in composition, yet have small variations both within and between the participants’ samples (Fig 4B).

**Figure 1:**
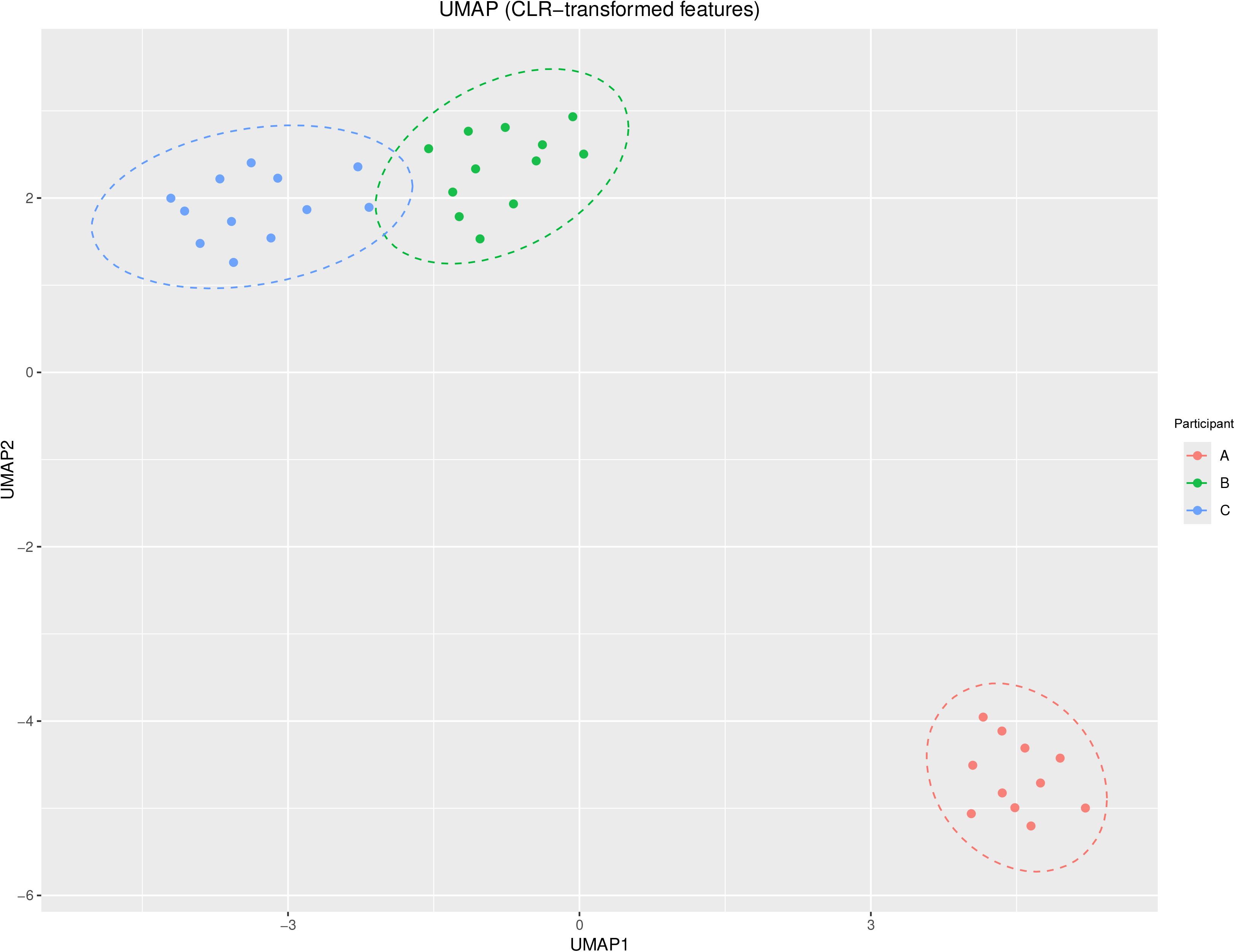
UMAP projecting the centered log ratio of the taxa counts in each participants’ urobiomes at each timepoint sequenced. Each point is a sample taken from the participants. Participants B and C associate with one another and have overlapping dispersions (dotted ellipses), while participant A maps away from both other participants. Ellipses were calculated using the stat_ellipse() function to fit a multivariate 95% confidence t-distribution across the centroids.

**Figure 2:**
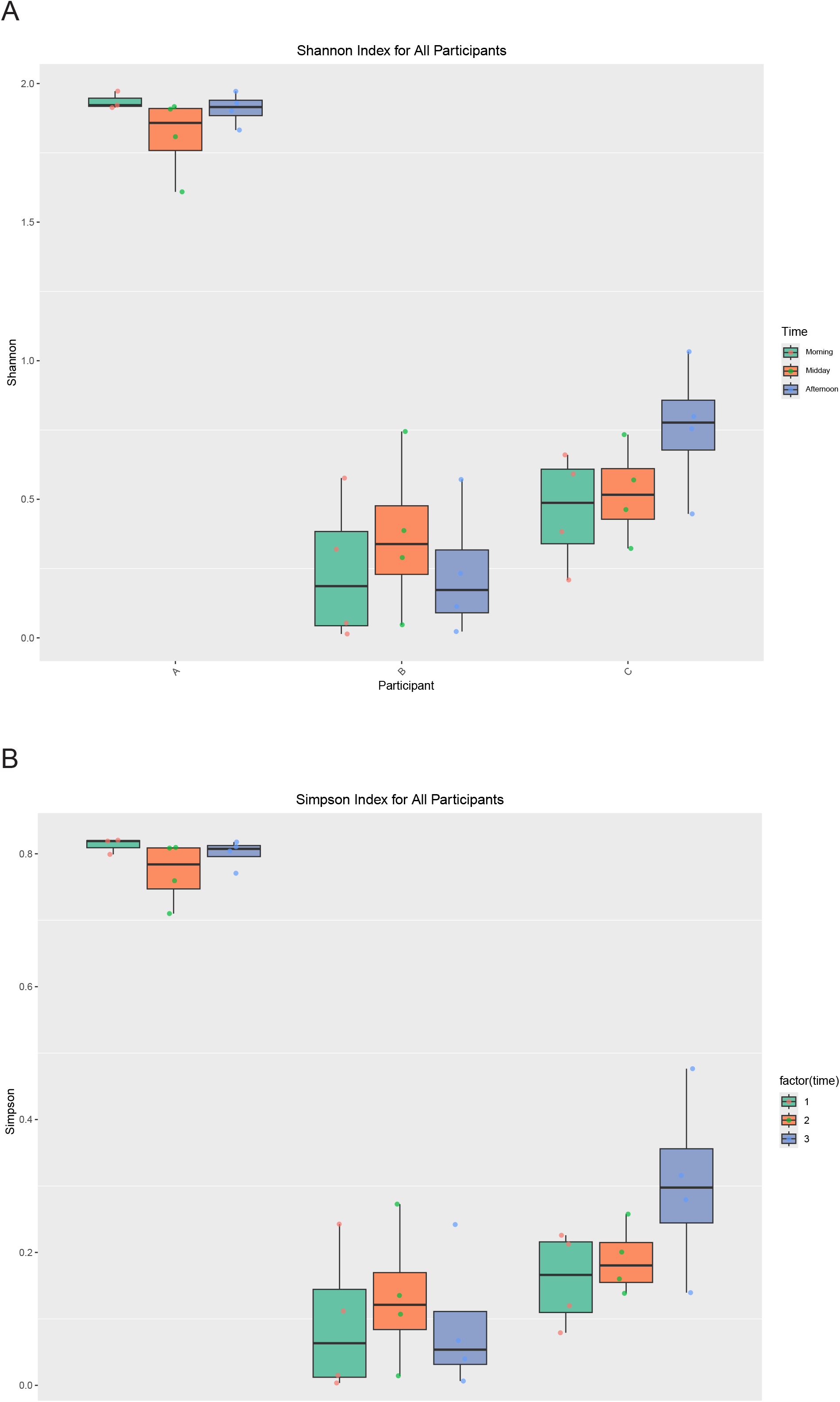
Alpha diversity indexes for each participant clustered based on the time of day the sample was taken. In both measures of alpha diversity, participants B & C cluster together, while participant A is much greater in value; however, all participants had similar values of alpha diversity between each timepoint (morning, midday, and afternoon) with no statistical difference between any two timepoints within the same participant. A) Shannon diversity indexes. B) Simpson diversity indexes.

**Figure 3:**
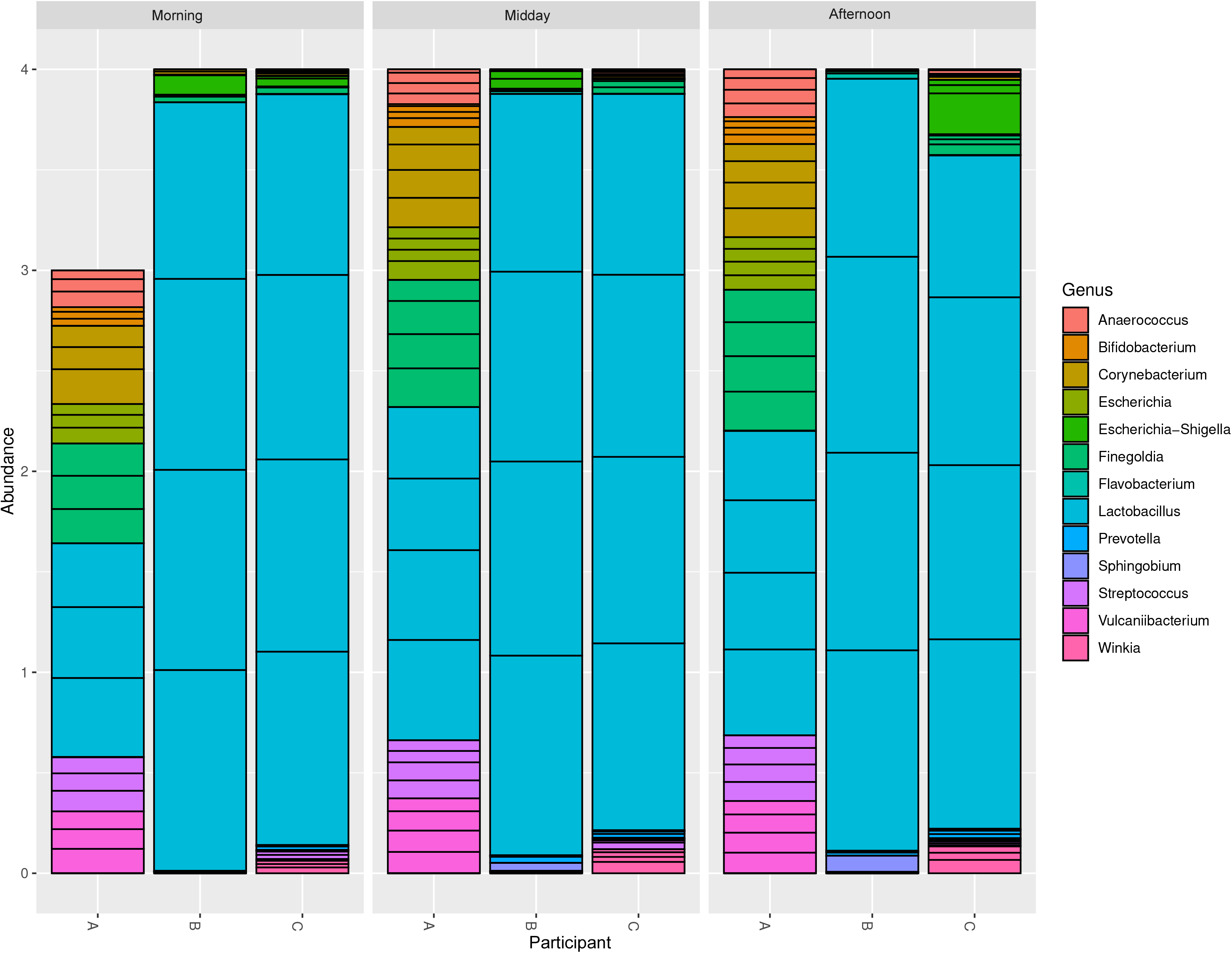
Bar plots displaying the spread of the top 20 Genera in the urines of three participants. Participants B and C are skewed towards species in the Genus *Lactobacillus*. Participant A had a more even distribution, yet only 10 Genera were confidently identified. Visually displaying the similarities between participants B and C.

**Figure 4:**
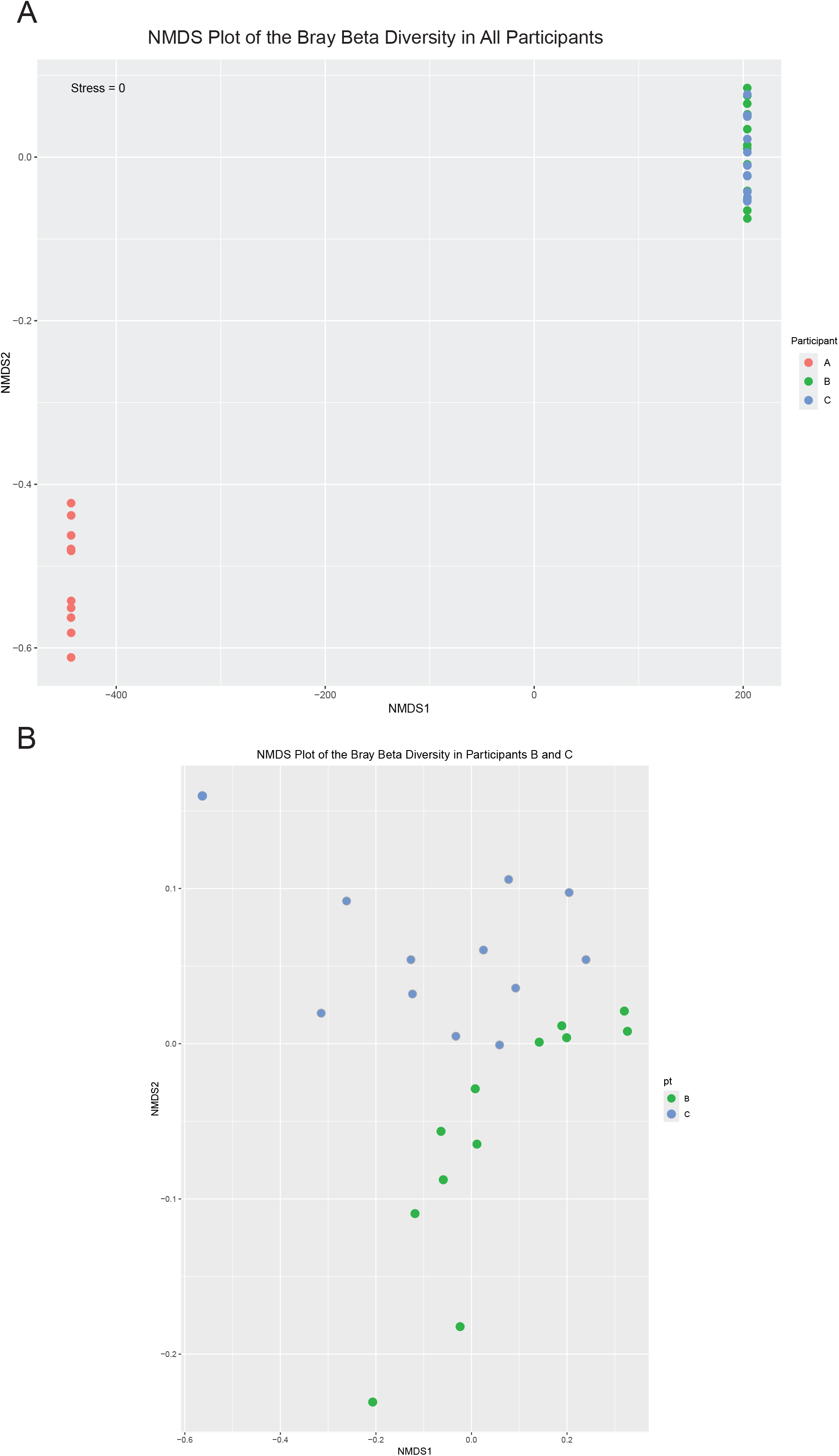
NMDS plots of the beta diversity indexes of each timepoint. A) NMDS plot for all participants. Participants B and C were highly similar, while participant A was highly dissimilar. B) To determine the differences between participants B and C, a plot excluding participant A was generated. Note that, due to the high similarity of the values, the scale is very narrow.

Taken together, the data suggest two different types of urobiomes: Urobiome type 1 consists of participant A with a high evenness yet a very low diversity, and Urobiome type 2 consists of participants B and C with a dominance of one Genus (*Lactobacillus*) and low diversity. Both patterns were maintained throughout each timepoint.

### Individual Analysis of the Urobiome Types

#### Urobiome Type 1

In the urobiome of participant A few bacterial 16S counts were observed; instead, more than 50% of the 16S counts mapped to the mitochondria (supplemental Fig 2). This correlated with the fact that the urines from participant A had more large particulates—assumed to be host cell sloughing—compared to urines from participants B and C. Removing the mitochondrial reads— shown in the results above and here—emphasizes the microbial counts, but some of the taxa are not known to be associated with humans in either commensal or pathogenic interactions (Fig 5). We present the data as it stands but interpret the results to ultimately suggest that participant A has a sparingly populated urobiome.

**Figure 5:**
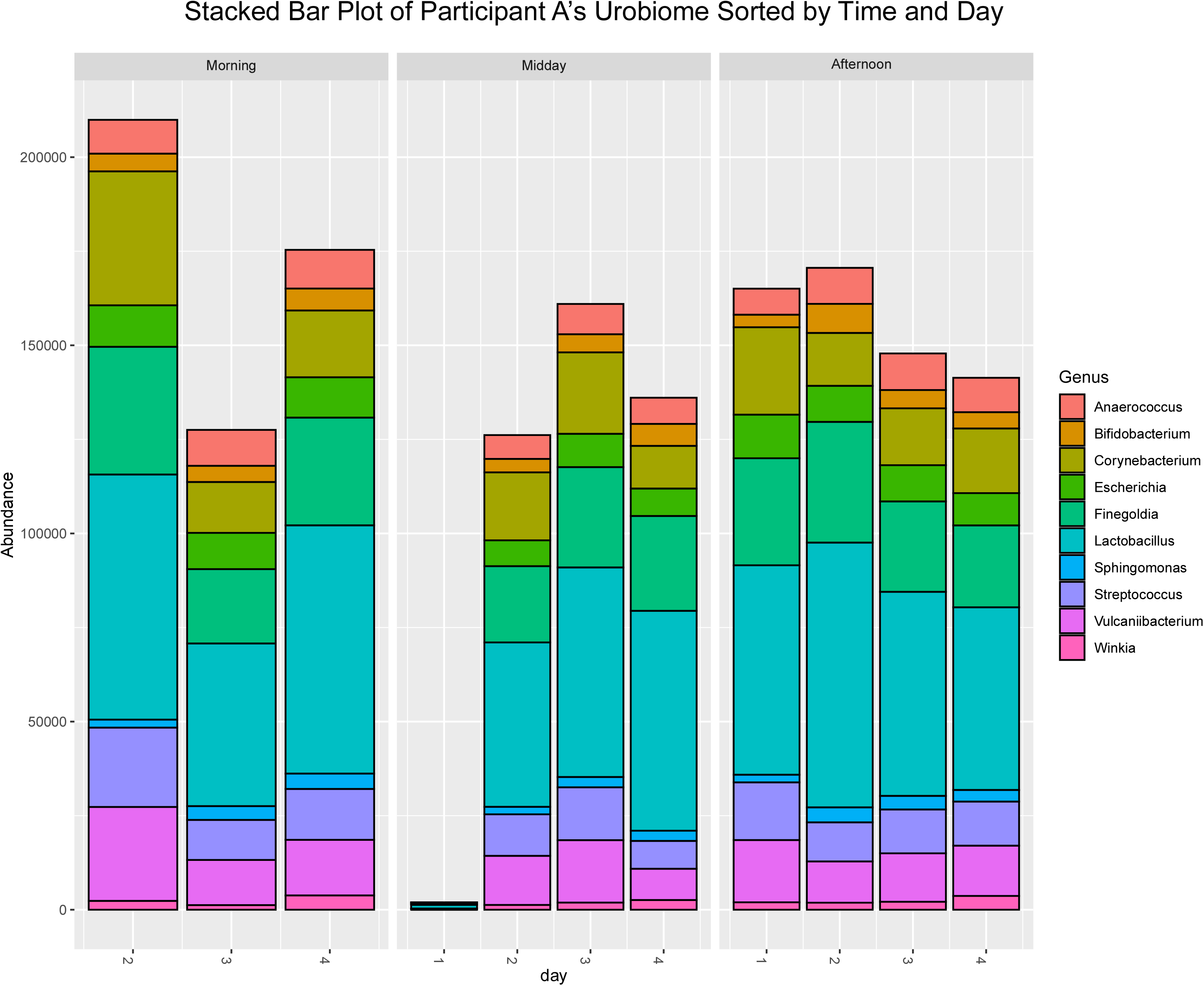
Stacked bar plots displaying the spread of the top 20 Genera in the urines of participant A. The urines of participant A were far more even in distribution of taxa than either of the other participants. There were only 10 Genera that were identified as being confidently present. The first midday timepoint had very little microbial presence, but the remaining days were nearly the same in distribution of taxa.

What was present consisted largely of *Lactobacilli* but counts of *Corynebacterium* and *Escherichia* were also present in high relative abundance. The remaining genera were mixed with microbes that are associated with the urinary tract *Streptococcus, Finegoldia, Anaerococcus, Winkia* and *Bifidobacterium* and microbes that are not associated with the urinary tract *Vulcaniibacterium*. In general, the total number of counts was greatly reduced compared to either of the other participants (10, vs 105 and 88). All Genus calls returned confidences of 100% except for *Escherichia* which returned confidences of greater than 60% (supplemental table 1).

#### Urobiome Type 2

Both participants B and C had a much less diverse, yet greater amount of sequence counts than participant A. These urobiomes both showed skews towards the genus *Lactobacillus* with minor fluctuations throughout the course of the experiment. Participant B had counts of *Escherichia* on six of the 12 timepoints (Fig 6A). Two of the time points also showed an increase in *Sphingobium* as well. Participant C had some calls of *Streptococcus* throughout the day, while a fraction of the afternoon readings on days 1, 2 and 4 called to *Escherichia* (Fig 6B). However, beyond those exceptions, all remaining time points looked nearly identical for both participants.

**Figure 6:**
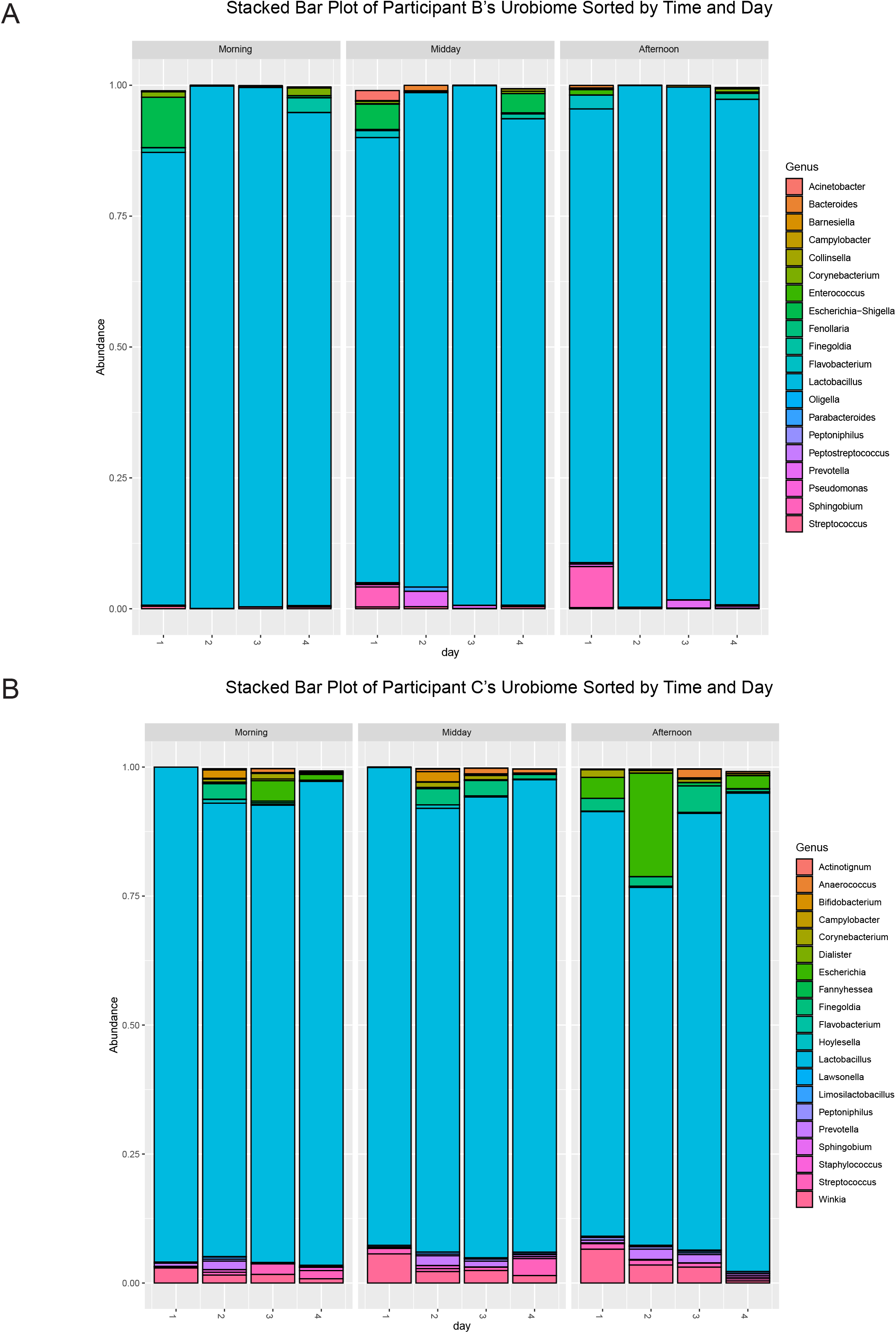
Stacked bar plots displaying the spread of the top 20 Genera in the urines of participants B and C. A) Stacked bar plot for participant B. The urines of participant B were skewed towards species in the Genus *Lactobacillus*. The first morning and midday timepoints had a spike of *Escherichia*, and the first midday and afternoon timepoints had a spike of *Streptococcus*. Otherwise, the urobiome is dominated by *Lactobacillus*. B) Stacked bar plot for participant C. The urines of participant C were skewed towards species in the Genus *Lactobacillus*. Participant C also had a constant presence of *Streptococcus* with only the final afternoon having a few counts of *Streptococcus*. The afternoon of day 2 also had a higher count of *Escherichia* than any other day; however, like participant B, the urobiome was skewed towards species in the Genus *Lactobacillus*.

All genus level calls returned 100% confidence (Supplemental Table 2 & 3). Species level analysis determined by BLCA indicated differences between the levels of *L. crispatus* and *L. inners*. Participant C especially displayed fluctuations in the levels of *L. inners* with a near 100% confidence at each *inners* call, while the *crispatus* calls returned confidences of approximately 34% (Supplemental Table 3).

## DISCUSSION

This is the first study to assess the daily variability of the urinary microbiome in post-menopausal females who never had a UTI. It is critically important to understand the natural urinary landscape of these participants to determine their protective mechanisms against invading uropathogens, as well as establish some baseline common findings to interpret the microbiome of females suffering from recurrent urinary tract infections, especially upon the completion of an antibiotic course as they return to “normal”.

We found two possible types of urobiomes: one with fewer and more evenly distributed genera, and another richer, less evenly distributed, and less comparatively diverse genera. We propose that the presence of *Lactobacillus* is protective for at least two of the participants, if not all three. Participant A and C’s urines were much less supportive of bacterial growth than participant B’s urines suggesting that nutrient limitation participates in protecting people from UTIs (9). Thus, protective bacterial taxa, nutrient availability, or both are factors that contribute to susceptibility.

This type of *Lactobacillus*-dominant urobiome has been noticed in healthy pre-menopausal females by a few other studies (3, 6). Price et al. 2020 tracked the urobiomes for 95 days and found that for the *Lactobacillus* dominant urobiomes little variation was seen throughout the study similar to what we report here (6). Price et al. also describes menstruation-dependent urobiome variability, though this effect was dependent on the individual (3). Unlike these studies, however, we calculate higher inverse Simpsons indexes and lower Shannon indexes which can be attributed to our lower overall number of participants. Our data implies that the urobiome patterns seen in younger females are maintained into postmenopausal age and may confer their protective effects into the later stages of life. Thus, understanding how the urobiome is influenced by daily activities, and hormonal fluctuations may impart protection against infection.

We speculate that multiple protection mechanisms exist. In the case of the diverse urobiomes (participant A), the diversity of their components may confer a protective effect against an impending pathogen. Like in the vaginal microenvironment, the catabolism of resources by lactobacilli may inhibit unestablished bacterial growth, and antibacterial products like L-phenyllactic acid may suppress infection establishment even further (18). Even if lactobacilli are not the protective agents, environmental and nutritional competition between an impending pathogen and other genera in a low nutrient environment coupled with frequent draining of the bladder may be sufficient to prevent infection in the early establishment phase (8, 9). In the case of the low diverse urobiome, we speculate that the increased epithelial cell turnover inhibits bacterial-host cell attachment limiting the perseverance of the bacteria in the bladder. If the bacteria cannot attach themselves to the bladder wall and effectively invade into the host cells, then the bacteria are more exposed to immune cells and toxins generated in the waste environment of the bladder and are thus more likely to be expelled during urination.

We acknowledge several limitations. First, 16S sequencing inherently introduces a bias towards those bacteria with 16S sequences that better align with the selected PCR primers, resulting in an artificial enrichment. Second, this study reports only bacterial components ignoring phage and fungi, both of which can also confer protection against infection (19, 20). Finally, clean-catch urine may introduce some non-urinary microbes into our analysis. Despite these limitations, we proceeded due to the cost effectiveness of 16S sequencing given that we had 36 samples to prepare and sequence, and that many studies have utilized 16S in the past giving us a large bank of data for comparison. Furthermore, since the participants were completely healthy (not admitted to the hospital) and provided samples multiple times a day over multiple days, we chose the clean-catch urine collection method. Clearly, in this context, catheterization, or cystocentesis urine collection methods would not have been ethically appropriate.

Overall, we present a limited but insightful analysis of the microbiomes of post-menopausal females with no history of UTI. Tracking the changes in these microbiomes over a two-week period with six timepoints per week indicated that the individual microbiomes are stable, at least in the short term, and trend towards an increase in the *Lactobacillus* genus when bacteria are present. One microbiome showed only an increase in mitochondrial 16S transcripts which we interpret as having a protective effect by increasing epithelial layer turnover inhibiting bacterial persistence in the bladder environment.

## Acknowledgements

The authors would like to thank the Genome Center at The University of Texas at Dallas for the services to support our research.

JH would like to thank Dr. Yeunhee Kim at the University of Texas at Dallas for her advice during the planning of the experimental process.

## Data Availability

Sequencing data has been made available through the NCBI Sequence Read Archive (SRA) at

